# Dynamic Processing of Displacement Loops During Recombinational DNA Repair

**DOI:** 10.1101/421990

**Authors:** Aurèle Piazza, Shanaya Shah, William Douglass Wright, Steven K. Gore, Romain Koszul, Wolf-Dietrich Heyer

## Abstract

Displacement-loops (D-loops) are pivotal intermediates of homologous recombination (HR), a universal DNA double strand break (DSB) repair pathway. We developed a versatile assay for the physical detection of D-loops *in vivo*, which enabled studying the kinetics of their formation and defining the network of D-loop formation and reversal pathways. Nascent D-loops are detected within 2 hrs of DSB formation and extended over the next 2 hrs in a system allowing break-induced replication. The majority of D-loops are disrupted in wild type cells by two pathways: one supported by the Srs2 helicase and the other by the Mph1 helicase and the Sgs1-Top3-Rmi1 helicase-topoisomerase complex. Both pathways operate without significant overlap and are delineated by the Rad54 paralog Rdh54 in an ATPase-independent fashion. This study uncovers a novel layer of HR control in cells relying on nascent D-loop dynamics, revealing unsuspected complexities, and identifying a surprising role for a conserved Rad54 paralog.

## Introduction

Homologous recombination (HR) repairs DNA double-strand breaks (DSBs) by exploiting an intact homologous double-strand DNA (dsDNA) molecule as a template. It proceeds in a succession of metastable intermediates, which confers flexibility and kinetic proof-reading at every non-covalent steps of the pathway (Heyer, 2015; Kanaar et al., 2008; Zinovyev et al., 2013). First, a helical filament of Rad51, a member of the RecA family, and associated proteins is assembled onto the 3’-protruding ssDNA generated upon DSB resection, which can be several kilobases long (Mimitou and Symington, 2008; Zhu et al., 2008). This multivalent filament harnesses the ssDNA sequence information to interrogate nearby dsDNA molecules (Bell and Kowalczykowski, 2016) as it dynamically weaves through the nuclear volume (Dion et al., 2012; Mine-Hattab and Rothstein, 2012). Upon successful identification of homology, Rad51/Rad54-catalyzed DNA strand invasion results in a nascent displacement loop (D-loop) intermediate of possibly varied architectures depending on length and whether the invasion involves the 3’-end or not (Wright and Heyer, 2014)and see below). The salient features of D-loops consist of at least a partially Rad51-free heteroduplex DNA (hDNA), a displaced ssDNA, and junctions at both extremities of the hDNA tract (**Fig. 1A**). Initiation of recombination-associated DNA synthesis by Polδ primed from the 3’-OH end of the invading strand represents a decision point in HR and is forming an extended D-loop. DNA synthesis restores the sequence information disrupted by the DSB. While disruption of nascent D-loops is a mechanism of anti-recombination, disruption of extended D-loops represent a mechanism of crossover avoidance enforcing non-crossover (NCO) outcome. Indeed, HR pathway choice (Gangloff et al., 2000; Shor et al., 2002), accuracy (Putnam and Kolodner, 2017) and outcome (Ira et al., 2003; Mazon and Symington, 2013; Mitchel et al., 2013; Prakash et al., 2009) are believed to at least partly rely on D-loop reversal (reviewed in refs. (Daley et al., 2014; Heyer, 2015)).

**Figure 1:**
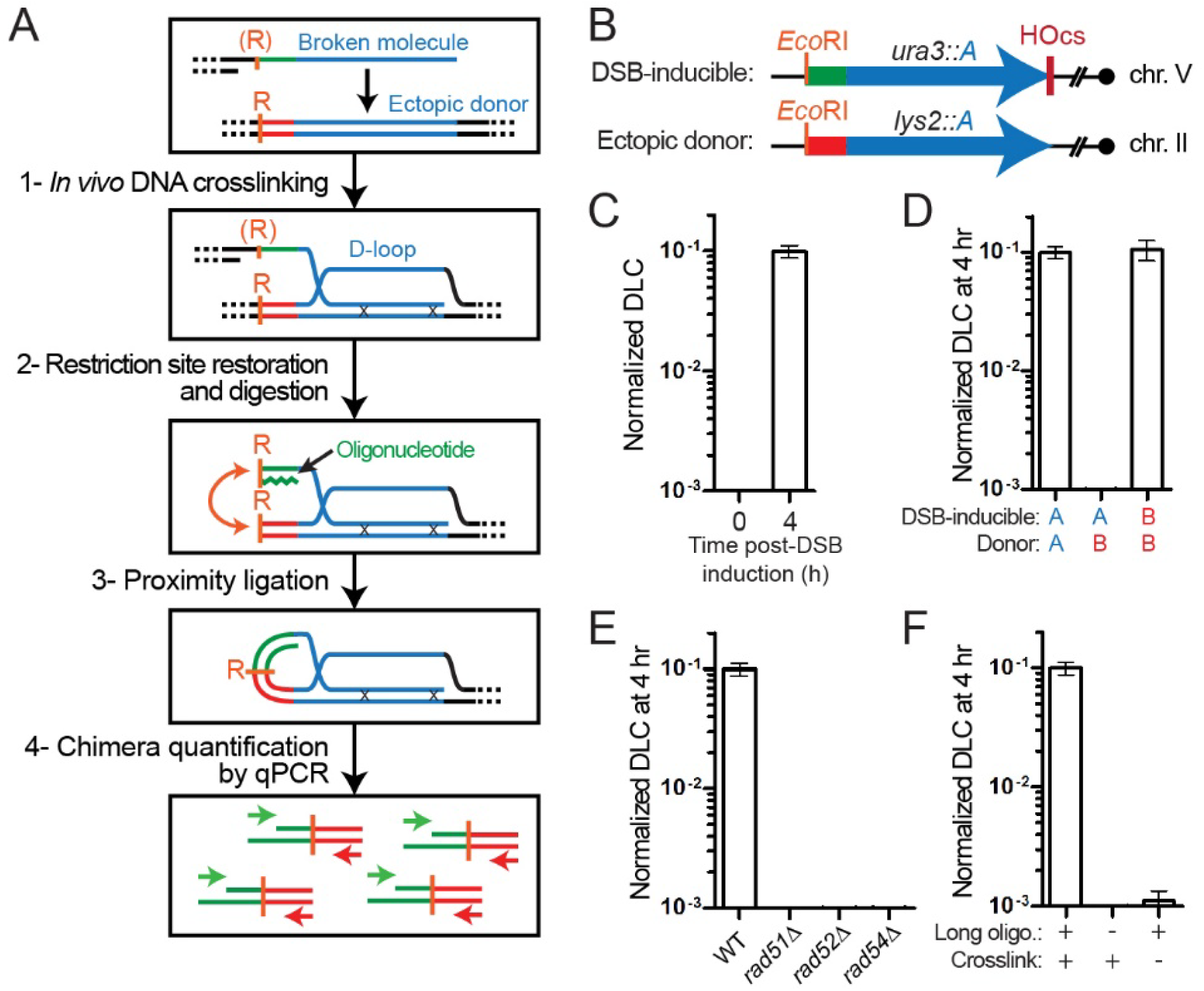
The DLC assay detects D-loops in *S. cerevisiae*. (A) Rationale of the DLC assay. (B) DSB-inducible construct and ectopic donor in haploid *S. cerevisiae*. The region of homology “A” is 2 kb-long. (C) DLC requires DSB induction on Chr. V. (D) DLC requires homology between the broken molecule and the donor. (E) DLC is HR-dependent. (F) DLC requires inter-strand DNA crosslink with Psoralen and restoration of the resected *Eco*RI site on the broken molecule. (G) Kinetics of DSB formation and DLC in wild type cells. (C-G) Bars represent mean ± SEM of at least a biological triplicate.

Several proteins have been implicated in joint molecule/D-loop turnover, whose defects specifically cause repeat-mediated genomic instability (Putnam et al., 2009). The 3’-5’ Srs2 helicase (putative homologs human FBH1 or RTEL1) is a negative regulator of HR likely acting at various steps of the pathway, Rad51 filament disruption and D-loop reversal. Srs2-deficient cells exhibit recombination-dependent lethality and genomic instability (Elango et al., 2017; Gangloff et al., 2000; Putnam et al., 2009). They also fail to mature NCO products in mitosis (Ira et al., 2003; Mitchel et al., 2013) as well as meiosis (Palladino, 1991), and are less prone to template switch during break-induced replication (BIR) (Ruiz et al., 2009), suggesting that Srs2 disrupts extended D-loops. *In vitro* Srs2 dismantles Rad51 filaments left unprotected by Rad55-Rad57 (Krejci et al., 2003; Liu et al., 2011; Veaute et al., 2003) and disrupts Rad51/Rad54-mediated nascent and extended D-loops (Liu et al., 2017). The 3’-5’ Mph1 helicase (human FANCM) funnels HR towards the NCO repair outcome (Mazon and Symington, 2013; Mitchel et al., 2013; Prakash et al., 2009; Sun et al., 2008; Tay et al., 2010), inhibits break-induced replication (BIR) (Jain et al., 2016; Luke-Glaser and Luke, 2012), and promotes template-switch during BIR (Stafa et al., 2014). Consistently, purified Mph1 disrupts short synthetic or Rad51/Rad54-mediated nascent and extended D-loops *in vitro* (Prakash et al., 2009; Sebesta et al., 2011; Zheng et al., 2011). The helicase/topoisomerase complex Sgs1-Top3-Rmi1 (STR, human BLM-TOPO3α-RMI1/2) limits formation and/or accumulation of various joint molecules including double Holliday Junction (dHJ) in somatic and meiotic cells (Bzymek et al., 2010; Kaur et al., 2015; Oh et al., 2007; Tang et al., 2015), inhibits CO and/or promote the NCO repair outcome of HR (Ira et al., 2003; Lo et al., 2006; Mazon and Symington, 2013; Mitchel et al., 2013; Tay et al., 2010), and inhibits BIR and long gap-repair (Jain et al., 2009; Jain et al., 2016). *In vitro*, STR dissolves dHJs (Cejka et al., 2010; Wu and Hickson, 2003) and D-loops (Fasching et al., 2015). dHJ dissolution requires both the Sgs1 helicase and Top3 topoisomerase activity. The STR complex is a complex D-loop disruption machine that can target protein-free D-loops through the helicase activity of Sgs1 and Rad51-Rad54-bound D-loops by the type 1A topoisomerase activity of Top3 (Fasching et al., 2015).

This large body of joint biochemical and genetic evidence established these factors as HR regulators enforcering the accuracy of the pathway (Heyer, 2015). Yet, the mechanisms by which they do so, including their precise substrates, interactions, and pathway organization remain elusive. Furthermore, the function of certain Rad51-ssDNA-associated proteins involved in regulating the DNA strand exchange reaction are not straightforwardly addressed *in vitro*, as the substrate and the conditions in which the reaction takes place are unknown. These limitations derive in good part from the technical inability to physically detect D-loops in somatic cells, unlike dHJ intermediates which can be detected by two-dimensional gel electrophoresis in meiotic and somatic cells (Bzymek et al., 2010; Schwacha and Kleckner, 1995). We developed the D-loop Capture (DLC) assay, a proximity ligation-based methodology that enables studying D-loop metabolism. This study confirms the D-loop disruption activities of multiple HR regulators and, by defining their interactions, reveals unanticipated complexities in nascent D-loops metabolism.

## Results

### The D-loop Capture (DLC) assay

The rationale of the DLC assay is depicted in **Fig. 1A**. The design is versatile, and we developed a specific protocol and controls for application in *Saccharomyces cerevisiae* (**Figs. 1B** and **S1**; **STAR Methods**). Because the donor lacks homology to the right side of the break, our genetic design purposedly restrict joint molecules to nascent and extended D-loops (**Fig. 1B**). The DSB-inducible and ectopic donor constructs are located at interstitial chromosomal loci and represent untethered and unconstrained location that have been extensively used by others (Agmon et al., 2013; Inbar and Kupiec, 1999; Mazon and Symington, 2013; Mine-Hattab and Rothstein, 2012). Following site-specific DSB induction (step 1) and DNA strand invasion at the donor, *in vivo* inter-strand DNA crosslinking with psoralen (Oh et al., 2009) covalently links the hybrid DNA (hDNA) within the D-loop to preserve its structure (step 2). The restriction site ablated by DSB resection is restored by annealing a long complementary oligonucleotide (step 3). Following restriction digestion (step 3), the ligation reaction performed in dilute conditions leads to preferential ligation of tethered DNA extremities, *i.e.* those held together by the crosslinked hDNA (step 4), a rationale common to all 3C-like approaches (Dekker et al., 2002). The unique chimeric ligation product is quantified by qPCR (step 5; details on normalization see **STAR Methods** and **Fig. S1**), and is referred to as the DLC signal.

As expected, the DLC signal depends on DSB formation (**Fig. 1C**), homology between the broken and donor molecules (**Fig. 1D**), and the central HR proteins required for filament assembly (Rad52) (Zelensky et al., 2014) and DNA strand invasion (Rad51 and Rad54) (Petukhova et al., 1998) (**Fig. 1E**). Contrary to previous proxy assays for DNA strand invasion (Rad51 ChIP at the donor locus)(Sugawara et al., 2003), the DLC signal requires homology (Renkawitz et al., 2013) and Rad54, consistent with biochemical data (Petukhova et al., 1998; Wright and Heyer, 2014). Together with the reliance of DLC on homology, D-loop stabilization by psoralen crosslinking and restoration of the restriction site eliminated by resection (**Fig. 1F**), these results demonstrate that the DLC assay detects D-loops and not nonspecific contacts between the broken and the donor molecule.

### Limitations of the DLC assay

A first limitation of the DLC assay resides in the psoralen-mediated inter-strand crosslink density (≈1/500 bp) (Oh et al., 2009). Since the *in vivo* hDNA length distribution is unknown, the DLC assay cannot distinguish between a single long and several shorter D-loops comprising the same total length of hDNA (**Fig. S2A**). Consequently, a change in DLC can reflect either an increase of the average hDNA length or an increase of the amount of D-loops in the cell population.

Second, upon long-range DNA synthesis, the D-loop will move away from the homologous donor loci and thus from the upstream restriction site. If it migrates past a downstream *Eco*RI site (located 11.1 kb away in our design), it will cause a physical uncoupling between the hDNA (*i.e.* the crosslink point between invading and donor molecules) and the upstream restriction site used for DLC chimera formation, thus precluding proximity ligation of both partners (**Fig. S2B**). In this study, we focus mainly on nascent D-loops, *i.e.* before extension by DNA polymerase, avoiding this potential limitation.

### Nascent and extended D-loops are temporally resolved

The fast and synchronous DSB induction upon HO expression (≈90% molecules cleaved within 30 minutes) enables kinetic study of the subsequent repair steps (Connolly et al., 1988; Hicks et al., 2011; White and Haber, 1990). D-loop formation is first detected 1 hr after DSB induction and increases 40-fold over 3-4 hr when it peaks before declining slightly (**Fig. 2A**). D-loop extension (monitored with another recently developed assay (Piazza et al., 2018); **Fig. S3**) follows with a ≈2 hrs delay: first detected at 4 hr, it peaks at 6 hr and plateaus (**Fig. 2A**). The mean time between half the DSB are formed (≈20 min) and half the maximum DLC and DLE signals are reached is DLC50 = 122 ± 13 min and DLE50 = 278 ± 23 min, respectively (**Fig. 2B**). This delay enables the separate study of nascent and extended D-loops at 2 and 6 hr post-DSB induction, respectively.

**Figure 2:**
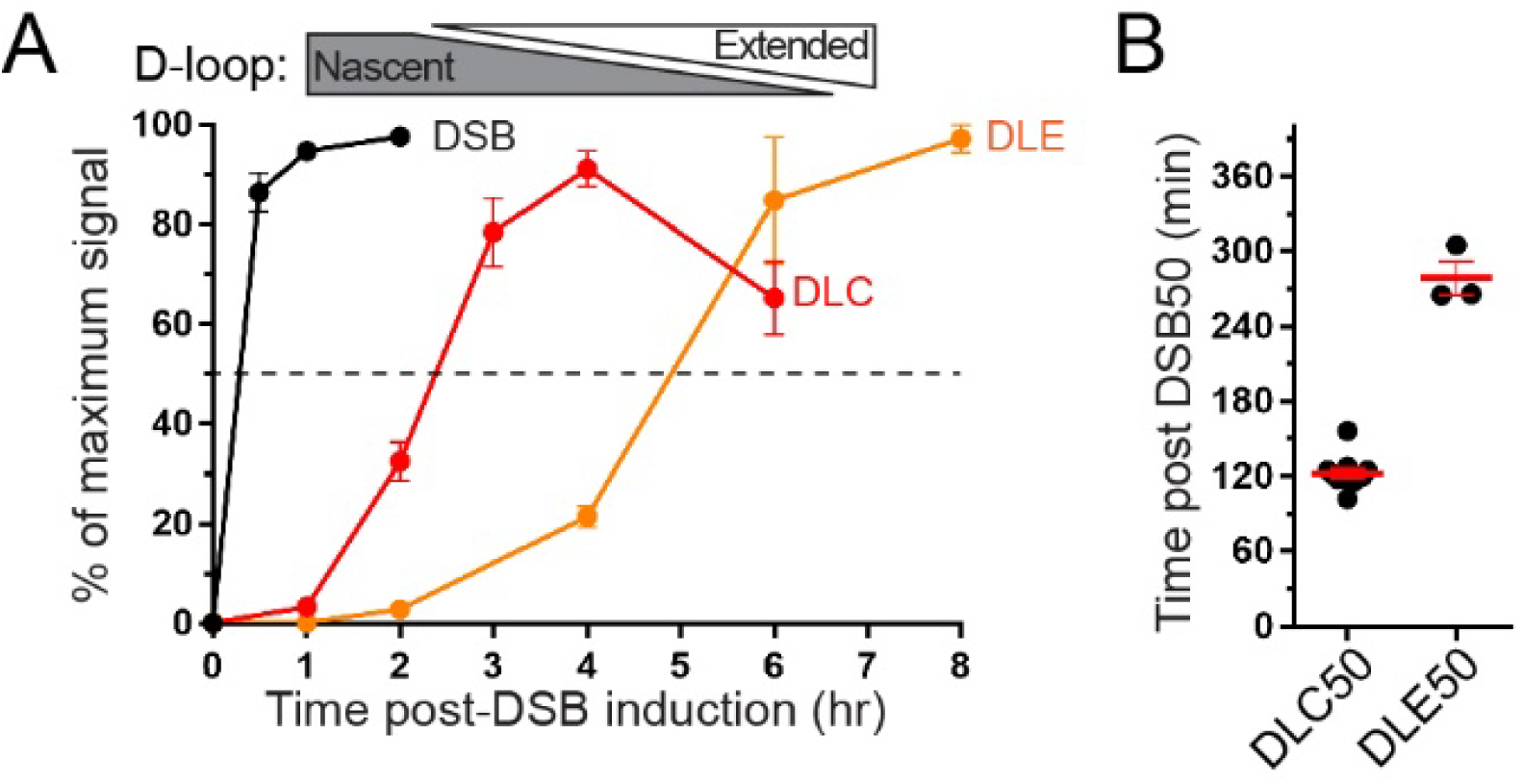
D-loop formation and extension kinetics. (A) Kinetics of DSB formation, DLC, and D-loop extension (DLE, see **Fig. S3**) (Piazza et al., 2018). DLC and DLE represent the mean ± SEM of 21 and 4 biological replicates, respectively. (B) DLC50 and DLE50 represent the time between 50 % of DSB formation and 50 % of the maximum DLC and DLE signal, respectively.

### Role of Srs2, Mph1 and STR in the dynamics of nascent and extended D-loops

We addressed the role of the Mph1 and Srs2 helicases as well as the helicase-topoisomerase STR complex in nascent and extended D-loop metabolism. Deletion of either *MPH1* or *SRS2* results in a significant 2-3 fold DLC increase at all time points (**Figs. 3A** and **3C**, respectively). The decrease observed between 2 and 4-6 hours in the *srs2Δ* mutant, from 2.9 to 1.8-fold over WT levels, is not statistically significant (**Fig. 3C**). The ATPase-deficient *mph1-D209N* and *srs2-K41A* mutants also exhibit elevated DLC levels compared to wild type, not significantly different from the corresponding deletion mutants (**Figs. 3B** and **3D**, respectively). Thus, both helicases inhibit the DLC signal, and hence steady-state levels of D-loops in wild type cells depend on their catalytic activity. These results are consistent with earlier biochemical evidence showing that Mph1 and Srs2 disrupt both Rad51/Rad54-made nascent and extended D-loops in an ATPase-dependent fashion (Liu et al., 2017; Prakash et al., 2009; Sebesta et al., 2011).

**Figure 3:**
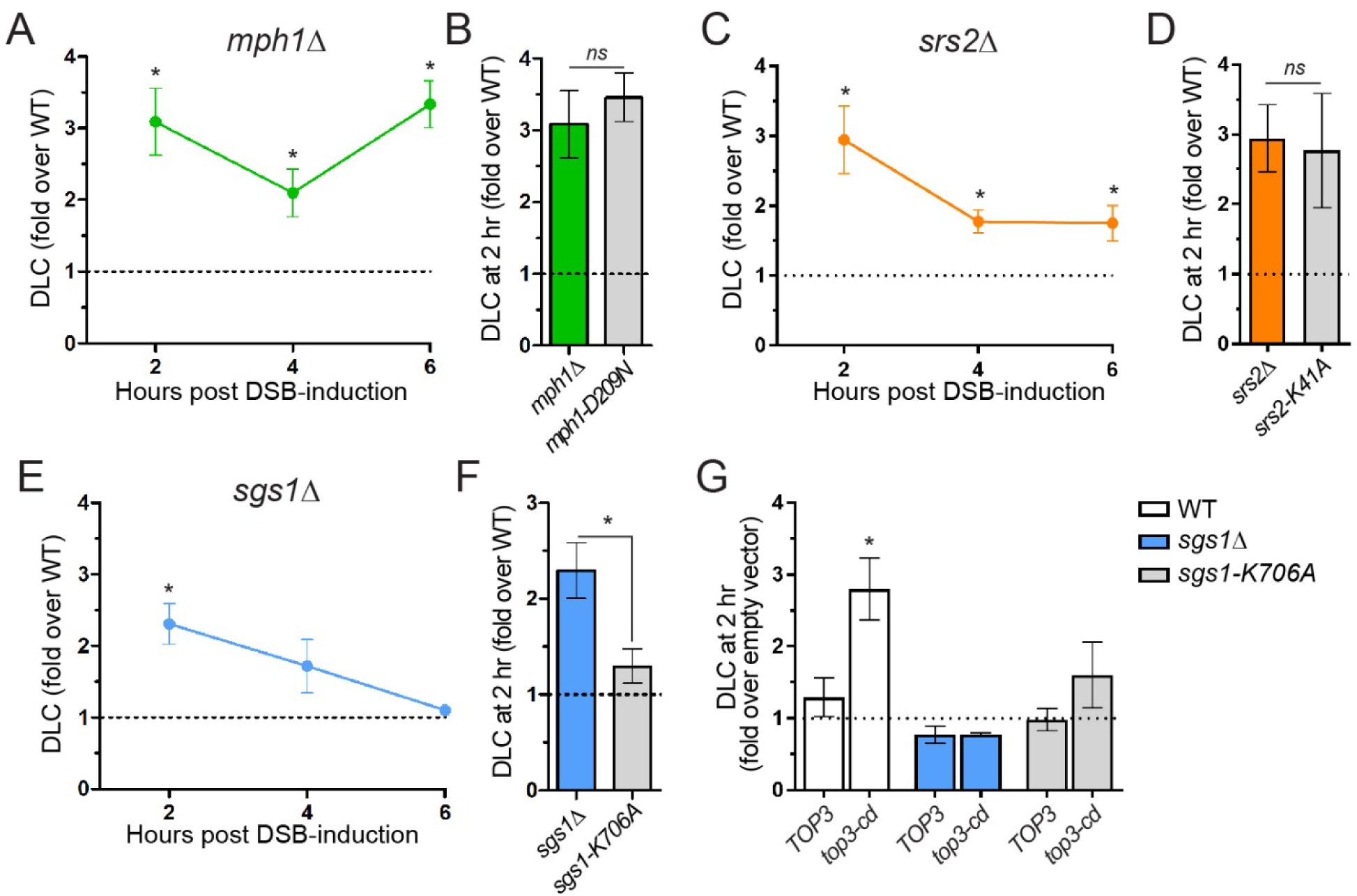
D-loop regulation by Mph1, Srs2 and STR. (A) Mph1 inhibits DLC at all time points. (B) DLC inhibition depends on the helicase activity of Mph1. (C) Srs2 inhibits DLC at all time points. (D) DLC inhibition depends on the helicase activity of Srs2. (E) Sgs1 inhibits DLC only 2 hr post-DSB induction and (F) in an ATPase-independent manner. (G) DLC inhibition by STR depends on Top3 catalytic activity. This inhibitory activity is epistatic to *SGS1* but independent of its helicase activity. Data represent mean ±SEM of at least biological triplicates. * indicates statistical significance (p<0.05). ns: not significant.

In contrast, STR significantly inhibits DLC only at the earliest time point: a *sgs1Δ* mutant exhibits a significant 2-fold DLC increase at 2 hrs post-DSB induction but not at 4 and 6 hrs (**Fig. 3E**). Unlike Mph1 and Srs2, DLC in the ATPase-deficient *sgs1-K706A* mutant is significantly lower than in the deletion mutant and not significantly different from wild type (**Fig. 3F**), indicating that the STR inhibitory effect requires the physical presence of Sgs1 but is largely independent of its helicase activity. Previous genetic observations showed that a subset of STR roles requires the physical presence of Sgs1 and its ability to interact with Top3, but is independent of its helicase activity (Jain et al., 2009; Lo et al., 2006; Weinstein and Rothstein, 2008). Furthermore, the topoisomerase activity of Top3, but not the helicase activity of Sgs1, is required for STR to disrupt Rad51/Rad54-mediated D-loops in reconstituted biochemical reactions (Fasching et al., 2015). To avoid working with the slow-growing and suppressor-prone *top3Δ* mutant, we addressed the role of the topoisomerase activity of Top3 upon induced overexpression of the dominant-negative catalytic-deficient *top3-Y356F* (referred to as *top3-cd*) mutant (Oakley et al., 2002). Overexpression of *TOP3* does not affect DLC, indicating that excess Top3 is unlikely partnered with Rmi and Sgs1 and ineffective at DLC inhibition (**Fig. 3G**). However, overexpression of *top3-cd* leads to a ≈2.5-fold DLC increase over the empty vector control (**Fig. 3G**), an increase similar to that observed in a *sgs1Δ* mutant. *TOP3* and *SGS1* are epistatic, as neither *TOP3* not *top3-cd* overexpression affects DLC levels in a *sgs1Δ* mutant (**Fig. 3G**). Overexpression of *top3-cd* in *sgs1-K706A* cells yields an intermediate, although non-significant, effect compared to either wild type or *sgs1Δ* cells (**Fig. 3G**). This intermediate effect may suggest a subtle contribution of Sgs1 helicase activity to STR-mediated D-loop disruption. These results show that nascent D-loop disruption requires the topoisomerase activity of Top3-Rmi1, and the physical presence but largely not the helicase activity of Sgs1. The non-catalytic role of Sgs1 could be to promote DNA strand passage during the Top3-Rmi1-mediated decatenation (Cejka et al., 2012), a reaction believed to underlie disruption of relatively short D-loops by STR (Fasching et al., 2015). Alternatively, Sgs1 may help target Top3-Rmi1 to its substrates.

In conclusion, these results provide direct evidence for three D-loop disruption activities in yeast. STR disrupts nascent D-loops in a topoisomerase-dependent and mostly helicase-independent fashion. Srs2 and Mph1 disrupt both nascent and extended D-loops in a helicase-dependent fashion.

### STR and Mph1 are part of the same nascent D-loop disruption pathway

We next investigated the genetic interactions between these D-loop disruption activities, focusing on nascent D-loop regulation. Cells defective for both Mph1 and STR do not exhibit additional DLC increase: the DLC profile in the *mph1Δ sgs1Δ* double mutant resembles that of a *sgs1Δ* single mutant at all time points (**Fig. 4A**). Furthermore, *top3-cd* overexpression in the absence of Mph1 does not yield further nascent DLC increase (**Fig. S4A**). This epistatic relationship is independent of the helicase activity of STR (**Fig. 4B**). These results indicate that STR and Mph1 operate in the same nascent D-loop disruption pathway. Regulation of extended D-loops is more complex, as the increased DLC observed at 6 hr in the *mph1Δ* mutant depends on STR (**Fig. 4A**), suggesting an antagonistic role in this context.

**Figure 4:**
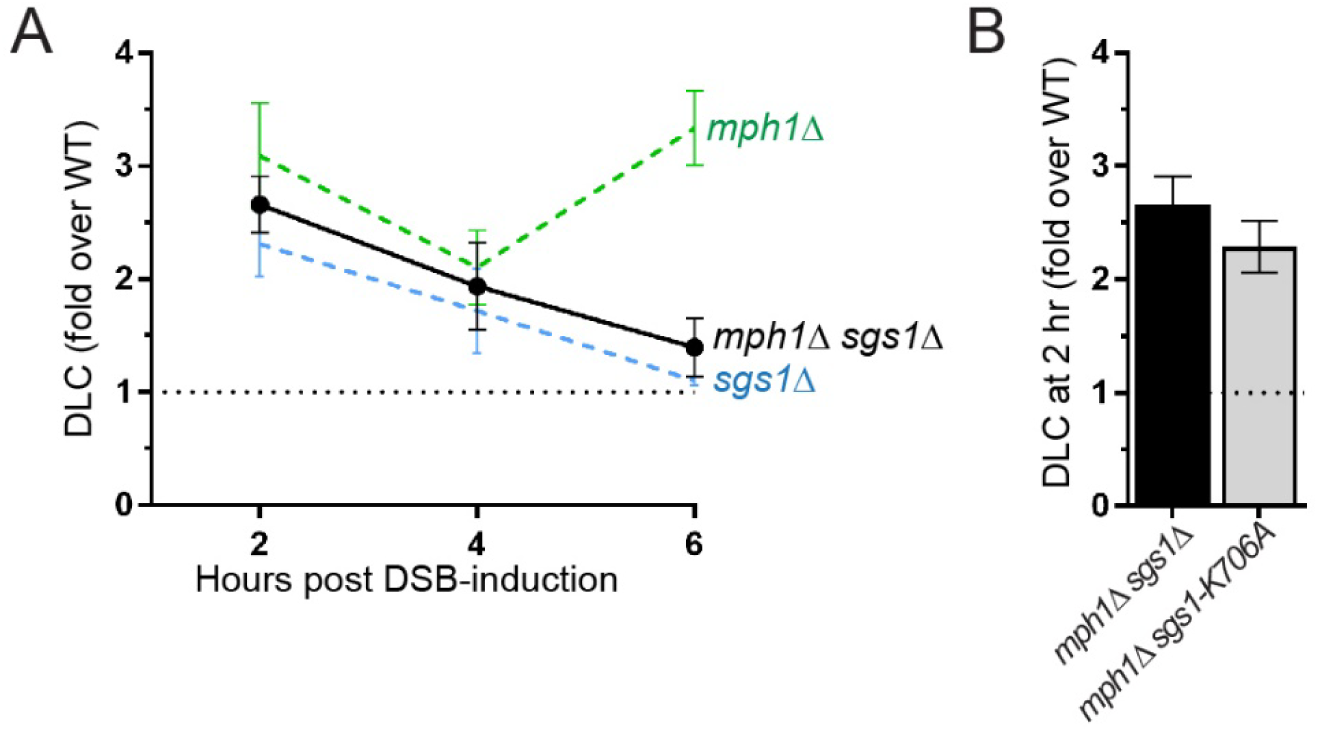
Mph1 and STR belong to the same D-loop disruption pathway. (A) *MPH1* and *SGS1* are epistatic in nascent DLC inhibition. (B) The helicase activity of Sgs1 plays no role in nascent D-loop processing in the absence of Mph1. Data represent mean ±SEM of at least biological triplicates. * indicates statistical significance (p<0.05).

### STR-Mph1 and Srs2 are independent pathways targeting non-overlapping nascent D-loop substrates

We next addressed the genetic interactions of *SRS2* with *STR* and *MPH1* on nascent D-loop metabolism. First, overexpression of *top3-cd* in a *srs2Δ* mutant causes a significant 1.9-fold DLC increase over the empty vector control (**Fig. 5A**), suggesting that Srs2 operates in a different D-loop disruption pathway than STR. In order to avoid the HR-dependent synthetic sickness of the double-deletion mutants (Gangloff et al., 2000; Lee et al., 1999; Mullen et al., 2001; Prakash et al., 2009) that are prone to acquire suppressors, we used an auxin-inducible degron version of Srs2 (Srs2-AID; **Methods**). Srs2-AID depletion induced prior to DSB formation (**Figs. 5B** and **S5D**) causes a 2.4-fold DLC increase, similar to that observed in a *srs2Δ* mutant (**Fig. 5C**). This result validates the *SRS2-AID* system for effectively depleting Srs2 protein and function. Srs2 degradation also leads to a significant DLC increase in the *sgs1Δ* and *mph1Δ* mutant background (**Fig. 5D**). Importantly, the *absolute* DLC increase observed upon Srs2 depletion in these mutants is not significantly different from what is seen in a wild type strain (**Fig. 5E**), indicating that the defect imparted by the absence of Srs2 is additive to that of the other mutations. The absence of epistasis or synergy but apparent additivity between the Srs2 and the STR-Mph1 disruption pathways indicate (i) that they target different nascent D-loop substrates, and (ii) that these substrates do not interconvert.

**Figure 5:**
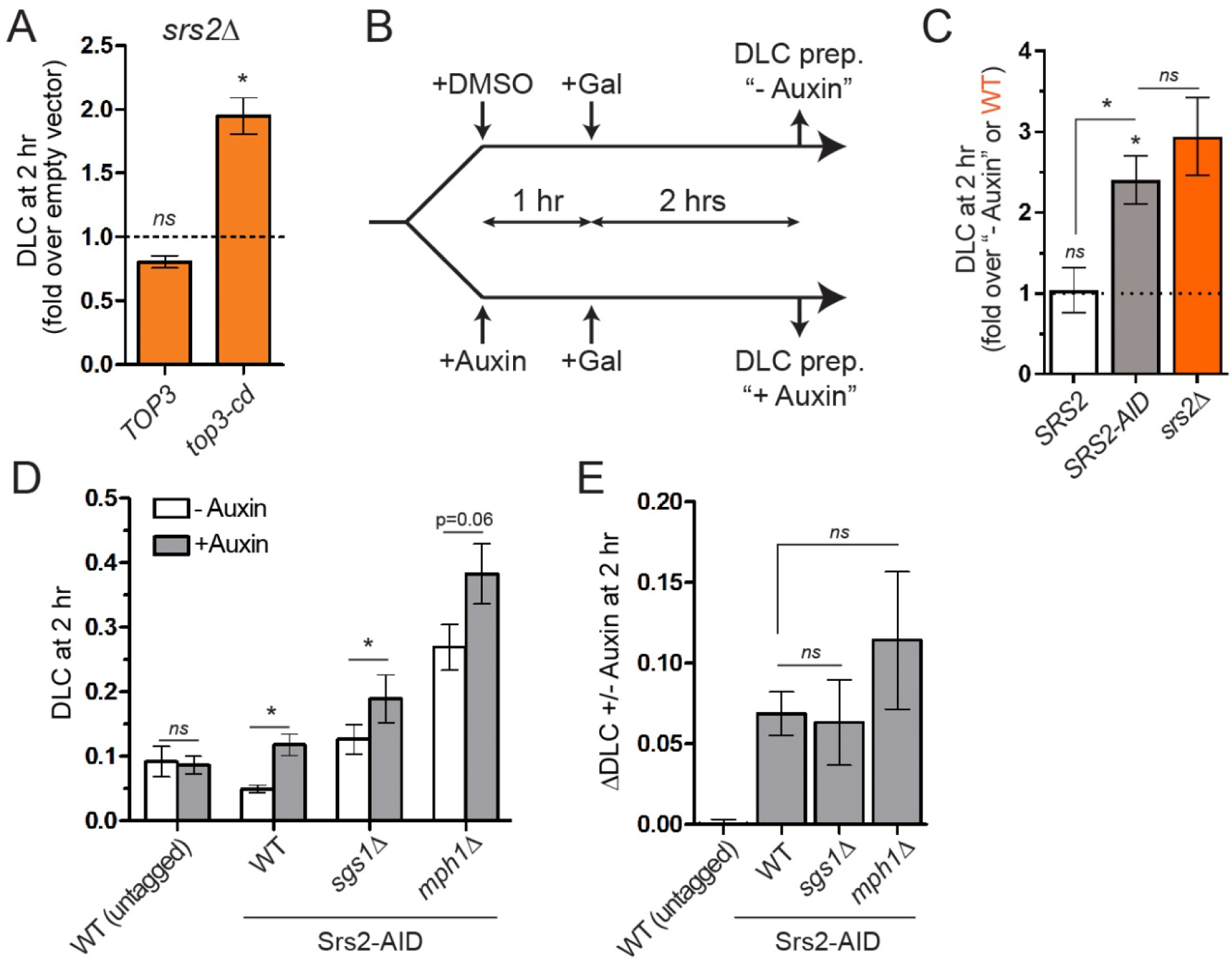
Srs2 and Mph1-STR are part of two non-overlapping nascent D-loop disruption pathways. (A) Overexpression of *top3-cd* in a *srs2Δ* mutant causes an additional nascent DLC increase. (B) Experimental scheme for Auxin-induced degradation of Srs2-AID. (C) Srs2-AID degradation upon auxin addition mimicks the *srs2Δ* mutant. (D) DLC increases in wild type, *sgs1Δ*, and *mph1Δ* strains upon Srs2-AID degradation. (E) Absolute extent of DLC increase in wild type, *sgs1Δ* and *mph1Δ* strains upon Srs2-AID degradation. Data represent mean ±SEM of at least biological triplicates. * indicates statistical significance (p<0.05). ns: not significant.

### Rdh54 inhibits nascent D-loops in an ATPase-independent fashion as part of the STR-Mph1 pathway

We sought to determine the apical determinant(s) of these two nascent D-loop disruption pathways. We surmised that it should involve components of the DNA strand invasion apparatus. Rdh54 (formerly known as Tid1) is a DNA translocase (Nimonkar et al., 2007; Prasad et al., 2007) primarily known for its role in meiosis (Klein, 1997; Shinohara et al., 1997), where it acts with Dmc1 for DNA strand invasion in a similar fashion as Rad54 acts with Rad51 in somatic cells (Nimonkar et al., 2012). Yet, contrary to its Dmc1 partner, Rdh54 is expressed in somatic cells (Lee et al., 2001), where it promotes inter-chromosomal template-switching during gene conversion and BIR (Anand et al., 2014; Tsaponina and Haber, 2014) and adaptation following DSB repair (Ferrari et al., 2013; Lee et al., 2001). Rdh54 physically interacts with Rad51 *in vitro* (Busygina et al., 2008; Petukhova et al., 2000) in an ATPase-independent fashion (Santa Maria et al., 2013) and in two-hybrids experiments in cells (Dresser et al., 1997). Furthermore, Rdh54 is recruited to DSBs in a Rad51-dependent fashion (Lisby et al., 2004), phosphorylated in response to DNA damage (Ferrari et al., 2013) and promotes engagement of dsDNA by Rad51 during homology search redundantly with Rad54 (Renkawitz et al., 2013). These observations suggest that Rdh54 is part of the Rad51-ssDNA filament in cells, similarly to Rad54 (Kiianitsa et al., 2006; Sanchez et al., 2013).

Deletion of *RDH54* causes a strong (≈3-fold) DLC increase at early time points corresponding to nascent D-loops (**Fig. 6A**). Intriguingly, the ATPase-defective *rdh54-K318R* mutant did not cause DLC increase, indicating that the early inhibitory effect of Rdh54 is exerted independent of its catalytic activity (**Fig. 6B**). *RDH54* is epistatic to *SGS1* and *MPH1*, as any mutant combination exhibits a similar ≈3-fold DLC increase over wild type (**Fig. 6C**). This result was corroborated upon *top3-cd* overexpression in *rdh54Δ* and *rdh54Δ sgs1Δ* cells, which did not exhibit significant increase over the empty vector control (**Fig. S6A**). Consequently, Rdh54 belongs to the STR-Mph1 nascent D-loop disruption pathway.

**Figure 6:**
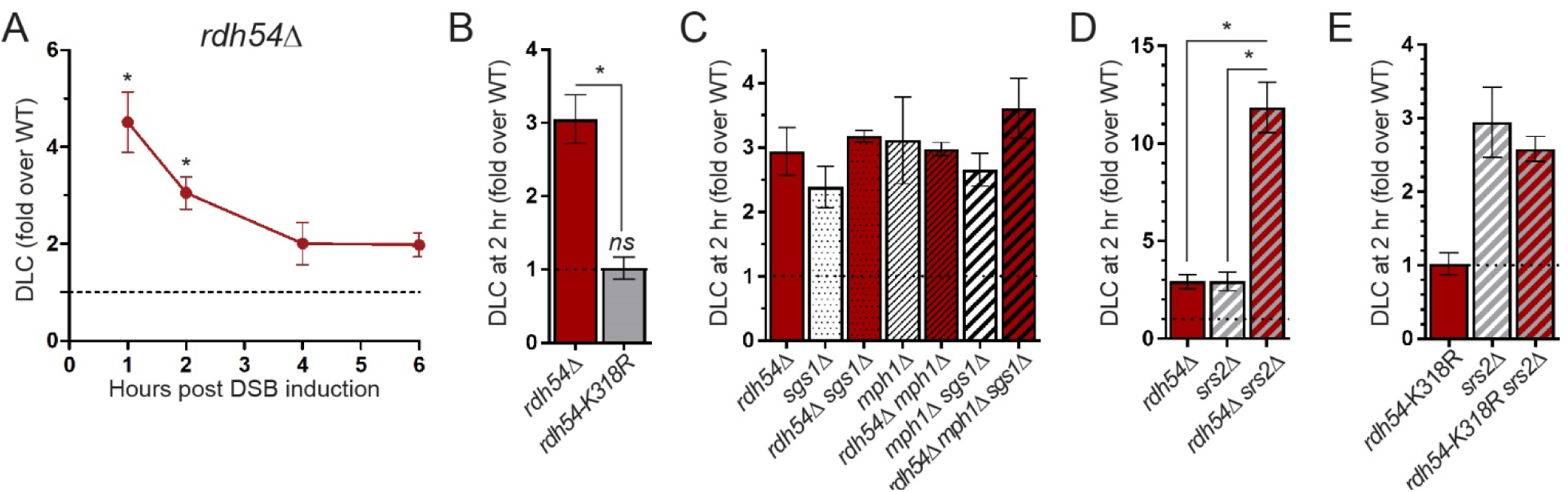
Rdh54 demarcates the two D-loop disruption pathways. (A) Rdh54 inhibits DLC at early time points. (B) Rdh54 inhibits DLC in an ATPase-independent fashion. (C) *RDH54* is epistatic to the STR-Mph1 nascent D-loop disruption axis. (D) Deletion of both *RDH54* and *SRS2* causes a synergistic DLC increase. (E) The ATPase activity of Rdh54 is not required for DLC inhibition in wild type cells or in a *srs2Δ* mutant. Data represent mean ± SEM of at least biological triplicates, except *rdh54-K318R srs2Δ* (biological duplicate). *p<0.05, paired t-test if compared to parallel wild type values. Unpaired otherwise.

### *RDH54* exhibits unique genetic interactions with *SRS2*

Combined defect of Srs2 with either STR or Mph1 causes HR-dependent synthetic sickness (Gangloff et al., 2000; Lee et al., 1999; Mullen et al., 2001; Prakash et al., 2009). Contrary to the *mph1Δ srs2Δ* and *sgs1Δ srs2Δ* mutants, *rdh54Δ srs2Δ* cells exhibit no major growth defect (**Fig. S6B**). Remarkably, the *rdh54Δ srs2Δ* mutant exhibits a strong nascent DLC increase, 12-fold over wild type levels (**Fig. 6D**). This increase is 4-fold higher than any single mutant, indicating that *SRS2* and *RDH54* synergistically inhibit nascent D-loop. Hence, Rdh54 is epistatic to the STR-Mph1 disruption axis but exhibits unique genetic interactions with the Srs2 pathway, both for DLC (synergy) and viability (no synthetic lethality). No synergy is observed in the *rdh54-K318R srs2Δ* mutant (**Fig. 6E**), confirming that the catalytic activity of Rdh54 is not required for its role in DLC inhibition. Based on these results, we propose that Rdh54 demarcates the two nascent D-loop disruption pathways (see below and **Figure 7**).

## Discussion

### The DLC assay: a versatile tool for studying DNA strand invasion in cells

The DLC assay enables physical detection in cells of D-loops, a central yet elusive intermediate of the HR pathway. More generally, this assay can detect any association between DNA molecules mediated by a regional hDNA crosslinkable with psoralen. The simple rationale related to 3C-type approaches and the large dynamic range (spanning over three orders of magnitude) of the qPCR readout grants applicability to experimental systems with less frequent DSB formation and/or DNA strand invasion than in our system. For example, we previously used a variant of this assay to detect the multi-invasion HR byproduct (Piazza et al., 2017). Since psoralen has already been extensively used in a variety of organisms (including mammalian cells) and DSB delivery by CRISPR-Cas9 is nearly universally applicable, we expect this versatile approach to open the way for physical study of the HR process in various organisms and cell types.

### Nascent D-loop dynamics: implications for HR fidelity and outcome

The steady-state DLC increase observed in various mutants reveals that the majority of D-loops formed at a perfectly homologous 2 kb-long donor are normally being disrupted in wild type cells. We provide direct evidence for three activities underlying this nascent and/or extended D-loop turnover in *S. cerevisiae* cells: Srs2 and Mph1 disrupt both nascent and extended D-loops in a helicase-dependent fashion, and STR disrupts nascent D-loops in a topoisomerase-dependent and largely helicase-independent fashion. Hence, D-loops exist in a dynamic equilibrium that is enforced by multiple regulatory activities, confirming prior conclusions drawn from end-point assays and modelling (Coic et al., 2011; Zinovyev et al., 2013). This dynamics is likely to account for the ∼2 hours delay observed between D-loop formation and extension.

While extended D-loop disruption is integral to the SDSA pathway, the role of disruption activities targeting perfectly homologous nascent D-loop is less straightforward within the current HR framework, but we speculate it participates in HR fidelity in three main ways. First, we showed previously that these activities inhibit multi-invasion-induced rearrangements, a tripartite recombination mechanism involving the cleavage of internal and terminal nascent D-loops (Piazza and Heyer, 2018; Piazza et al., 2017). It also applies to single D-loop cleavage causing half-CO (Deem et al., 2008; Hum and Jinks-Robertson, 2018; Mazon and Symington, 2013; Pardo and Aguilera, 2012; Smith et al., 2009). Second, nascent D-loop turnover is expected to enforce homology search stringency (and thus HR fidelity) by requiring several rounds of donor interrogation prior to initiation of recombinational DNA synthesis (Coic et al., 2011). Hence, a blind sampling engine that has no knowledge whether it has reached the best genomic target is afforded a “second chance”. Given that homology length stimulate and that sequence divergence inhibits sequence recognition by the Rad51-ssDNA filament (Bell and Kowalczykowski, 2016), this kinetic proofreading strategy is expected to funnel the searching molecule towards the longest and most similar donor available. Indeed, in the absence of a homologous donor to compete with a homeologous one, disruption cycles are futile and uncover the tolerance to mispaired bases of the Rad51-ssDNA homology search engine (Anand et al., 2017; Lee et al., 2017). Consequently, when studying the homology search process and its accuracy in cells, one has to consider not only the imperfectly stringent dsDNA sampling activity by the core Rad51-ssDNA filament, but also the context of multiple cycles of invasion/disruption. Third, owing to the extended D-loop turnover that requires to re-initiate the whole pathway, such a mechanism would disproportionately inhibit repair requiring long extension tracts, thus overpowering mutagenic repair such as BIR with invasion/(synthesis/)disruption cycles. Consistently, the three nascent D-loop disruption activities defined here (Mph1, STR, and Srs2) suppress BIR or long gap repair (Jain et al., 2009; Jain et al., 2016; Luke-Glaser and Luke, 2012; Ruiz et al., 2009; Stafa et al., 2014), even though we show that STR does not promote extended D-loop disruption.

Furthermore, the CO outcome of HR has been associated with longer gene conversion tracts than NCO outcome (Aguilera and Klein, 1989; Ahn and Livingston, 1986; Guo et al., 2017; Maloisel et al., 2004). Since COs pose a risk of LOH at the subsequent cell generation, they are suppressed in somatic cells. Cells impaired for STR and Mph1 activities are proficient in chromosomal DSB repair but are partially deficient in CO suppression (Ira et al., 2003; Mazon and Symington, 2013; Prakash et al., 2009). Since the CO outcome of HR likely entails the formation of a dHJ intermediate (Szostak et al., 1983), the CO suppression defect of these mutants has been proposed to result from both the avoidance of dHJ intermediates by disrupting extended D-loops by Mph1 (Crismani et al., 2012; Lorenz et al., 2012; Mazon and Symington, 2013; Prakash et al., 2009) as well as the dissolution of dHJ intermediates by STR to prevent their endonuclease-mediated resolution into CO (Bizard and Hickson, 2014). An expectation of such sequential activities is a synergistic increase in CO upon inactivation of both pathways. Yet, the combined deletion of *MPH1* and *SGS1* is not synergic nor even additive for CO formation (Mazon and Symington, 2013; Mitchel et al., 2013; Tay et al., 2010). It rather indicates that STR and Mph1 mainly suppress CO as part of the same pathway, congruent with our nascent D-loop disruption results. Since donor invasion by both ends of the DSB and/or a second-end capture of the displaced strand of the extended D-loop is required for dHJ and CO formation, it suggests that the nascent D-loop turnover supported by STR-Mph1 could be a component of the negative regulation of CO formation. The fact that nascent D-loop reversal participates of CO suppression might help explain the overlap between *cis* and *trans* determinants that protect both against ectopic recombination and CO, such as sequence divergence (Tay et al., 2010; Welz-Voegele and Jinks-Robertson, 2008), homology length (Inbar et al., 2000; Jinks-Robertson et al., 1993), and the aforementioned D-loop disruption activities (Myung et al., 2001; Putnam et al., 2009; Putnam et al., 2016). Hence, the fate of the HR outcome could be tied at least partly to the nature of the initial or early DNA joint molecule intermediate. This interpretation holds implications for the yet unresolved CO designation step during meiosis that has been proposed to occur at the level of an early and undetected HR intermediate (presumably a nascent or short extended D-loop) (Hunter and Kleckner, 2001). The mechanism could involve specific stabilization of a nascent or short extended D-loop intermediate to provide the time for a second invasion or end capture and dHJ formation.

### A complex network of proteins enforces D-loop dynamics

Genetic interaction between these regulatory activities also revealed unexpected complexities in nascent D-loop metabolism (**Figs. 4** and **5**). Indeed, STR with Mph1 and Srs2 define two independent nascent D-loop disruption pathways (**Fig. 7**). Importantly, combined elimination of any member of both pathways results in an additive increase in D-loop levels. This additivity implies that the two pathways target non-overlapping and non-interconvertible nascent D-loop species. Congruent with this idea, Mph1 and Srs2 defects behave differently with respect to HR outcome: Mph1 promotes NCO formation from substrates that can otherwise be salvaged as CO, while Srs2 substrates cannot and remain unrepaired in Srs2-defective cells (Ira et al., 2003; Mazon and Symington, 2013; Prakash et al., 2009).

**Figure 7:**
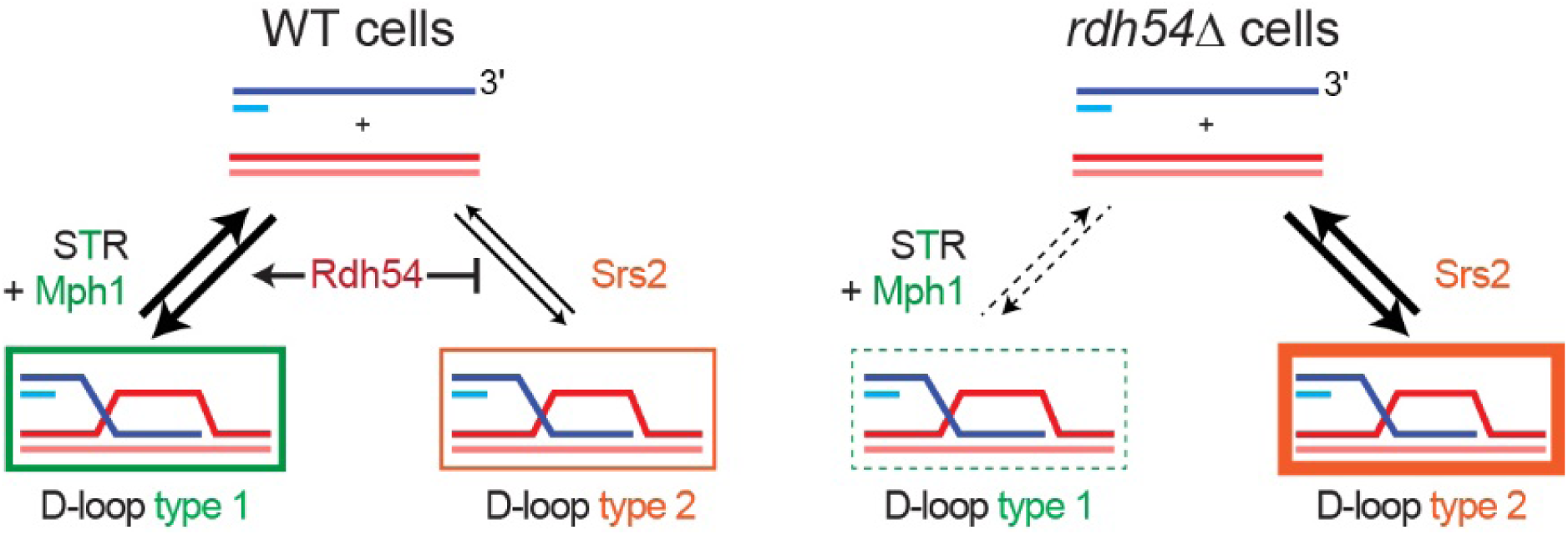
Model for the regulation the nascent D-loops by Rdh54, STR-Mph1 and Srs2. The box thickness schematizes DLC levels. Type 2 D-loops are possibly longer than Type 1, thus contributing more to DLC. For more details see **Fig. S6A**.

We propose a model for nascent D-loop metabolism by STR-Mph1 and Srs2 in which Rdh54 acts as a gate keeper in delineating the two disruption pathways (**Fig. 7**). Rdh54 promotes the formation of a substrate for the STR-Mph1 axis, while escapers are exclusively disrupted by Srs2. In particular, this model explains why, despite being part of the Mph1-STR D-loop disruption axis, defects of Rdh54 could synergize with those of a Srs2-defective strains in term of DLC (see below and **Fig. S6A** for more details).

How does Rdh54 exerts its demarcation role? And what distinguishes the two disruption pathways? The ATPase-independent role of Rdh54 rules out chromatin remodeling (Kwon et al., 2008), change in DNA supercoiling (Chi et al., 2006; Petukhova et al., 2000; Prasad et al., 2007), joint molecule disruption (Nimonkar et al., 2007), or Rad51 stripping from dsDNA (Chi et al., 2006) in the pathway demarcation process. Furthermore, the synergistic DLC increase specifically observed in *rdh54Δ srs2Δ*, but not in cells deficient for Srs2 and any member of the Mph1-STR disruption pathway, suggests that Srs2 substrates potentially contain longer hDNA than the substrates funneled by Rdh54 to the STR-Mph1 disruption pathway. Rdh54 could limit hDNA length by acting as a roadblock for Rad54-mediated hDNA formation (Wright and Heyer, 2014). Whether the hDNA length *per se* or architectural features (*e.g.* internal *versus* end invasion) distinguishes the two nascent D-loop species, and hence their targeting by one or the other pathway, remains to be addressed (**Fig. S6B**). Indeed, DNA strand invasion can occur not only at the 3’-OH extremity but also internally, although slightly less efficiently (Adzuma, 1992; Piazza et al., 2017; Wright and Heyer, 2014). Internal invasions likely exhibit altered architecture compared to terminal D-loop, both at the level of hDNA and the 5’ and 3’ junctions (Wright et al., 2018). How these structural features are recognized and guide differential processing by the Srs2 and STR-Mph1 (and possibly other) pathways remains to be addressed.

In conclusion, this work reveals a novel layer of HR control at the DNA strand invasion and nascent joint molecule levels, clarifies the interactions between several HR regulators, and suggests unanticipated roles for conserved Rad51-associated factors such as the Rad54 paralog, proteins of as yet poorly defined function.

## Acknowledgements

We thank members of the Heyer, Hunter and Kowalczykowski laboratories for helpful discussions, Ian Hickson for providing the *TOP3* overexpression vectors, and Hannah Klein for providing the *rdh54-K318R* mutant. We thank Abou Ibrahim-Biangoro and Noelle Cabral for help with strains construction. We are particularly grateful to Sue Jinks-Robertson and Pallavi Rajput for their critical reading of the manuscript.

## Authors contributions

AP and WDH conceived the project and wrote the manuscript, with input and editing from WW and RK. AP designed, performed and analyzed the experiments. SS ans SKG performed and analyzed the experiments.

## Methods

### Strains

The genotype of the haploid *Saccharomyces cerevisiae* strains (W303 background) used in this study are listed **Table S1**. They contain a copy of the *HO* endonuclease gene under the control of the *GAL1/10* promoter at the *TRP1* locus on Chr. IV (Pannunzio et al., 2008). A point mutation inactivates the HO cut-site at the mating-type locus (*MAT*) on Chr. III (*MAT***a-**inc or *MAT*α-inc). The heterozygous DSB-inducible construct replaces *URA3* on Chr. V (−16 to +855 from the start codon). The DSB-inducible construct contains the 117 bp HO cut-site(Fishman Lobell and Haber, 1992), a sequence “A” (+4 to +2068 of the *LYS2* gene), and a 327 bp fragment of the PhiX genome flanked by multiple restriction sites. The “A” donor replaces the *LYS2* gene on Chr. II. The *URA3* locus on Chr. V and of the *LYS2* locus on Chr. II have been chosen because of their interstitial and untethered location (Agmon et al., 2013; Berger et al., 2008; Duan et al., 2010; Mine-Hattab and Rothstein, 2012; Zimmer and Fabre, 2011), which represents a maximally demanding homology search situation extensively used by others to study ectopic HR repair (Agmon et al., 2013; Inbar and Kupiec, 1999; Mazon and Symington, 2013; Mine-Hattab and Rothstein, 2012). Since the region of homology to the ectopic donor “A” does not encompass the DSB site, this system prevents formation of later intermediates involving both ends of the DSB, so as to focus our study on the regulation of D-loop intermediates. BIR (the only available repair pathway) is discouraged by the presence of the centromere(Morrow et al., 1997) and is in any case lethal. This system thus prevents resumption of growth and invasion of the population by cells undergoing early repair. We showed previously that the formation of BIR product are not detected prior to 8 hrs after DSB induction (Piazza et al., 2018). The annotated sequences of the DSB-inducible and donor constructs are available as ape files in **Dataset S1**.

To investigate the genetic interaction of *SRS2* with *SGS1* and *MPH1* we used a conditional protein degradation system induced by auxin (Morawska and Ulrich, 2013). Srs2 is fused to its C-terminus to an auxin-inducible degron (AID) tag together with 9-Myc (referred to as Srs2-AID). The Srs2 Strains bearing the un-induced Srs2-AID construct and lacking either the *SGS1* or *MPH1* genes grow normally (**Fig. S4B**), indicating that the AID tag does not impair the essential Srs2 function in these mutants. The gene encoding the AID-specific E3 ubiquitin ligase OsTIR1 under the control of *pADH1* promoter is constitutively expressed from a centromeric vector (pRS314)(Sikorski and Hieter, 1989). Auxin (Sigma I5148) was dissolved in DMSO at 285 mM. In the absence of auxin (equivalent DMSO concentration), Srs2-AID appears slightly more potent in DLC inhibition than untagged Srs2 (**Figs. S4C** and **5D**). Srs2 is maximally depleted within 1 hr following auxin addition (2 mM final concentration) (**Fig. S4D**). Consequently Srs2-AID degradation is induced 1 hr before DSB induction (**Fig. 5B**). Auxin did not induce change of DLC, as shown in a treated strain bearing an untagged version of Srs2 (**Figs. 5C** and **5D**). The empty, *TOP3* and *top3-Y356F* over-expression vectors were kindly; provided by Ian Hickson (Oakley and Hickson, 2002). The genes are under the control of a pGAL1 promoter on a 2-micron multi-copy plasmid (pYES2). The *rdh54-K318R* (*rdh54-K352R* in the S288c reference) mutant was kindly provided by Hannah Klein (Chi et al., 2006). Other mutants were generated by traditional gene replacement with antibiotic-resistance or prototrophic genes by regular lithium-acetate transformation.

### Culture media

Synthetic dropout and rich YPD (1% yeast extract, 2% peptone, 2% dextrose) solid and liquid media have been prepared according to standard protocols(Treco and Lundblad, 2001). Liquid YEP-lactate (1% yeast extract, 2% peptone, 2% Lactate), Lactate-URA and Lactate-TRP (0.17% Yeast Nitrogen Base, 0.5% Ammonium Sulfate, respective 0.2% amino acids dropout, 2% Lactate) were made using 60% Sodium DL-lactate syrup. All the cultures were performed at 30°C.

### DLC assay

Cells were cultured to exponential phase in YEP-lactate and DSB at the HOcs on Chr. V was induced by HO expression upon galactose addition as described in (Piazza et al., 2017). A total of 2.10^8^ cells were collected before, or at various time post-DSB induction by galactose addition, pelleted and re-suspended in 2.5 mL of a Psoralen crosslinking solution (0.5 mg/mL Trioxsalen (Sigma-Aldrich T6137), 50 mM Tris HCl pH 8.0, 50 mM EDTA, 20% ethanol). Crosslink of cells was performed in a 60 mm petri dish upon long wave (365 nm) UV irradiation in a Spectrolinker XL-1500 (Spectroline) for 15 minutes with permanent orbital agitation (50-70 rpm). Cells were washed in 50 mM Tris HCl pH 8, 50 mM EDTA and the pellet stored at −20°C. Cells were spheroplasted in a zymolyase solution (0.4 M Sorbitol, 0.4 M KCl, 0.5 mM MgCl2, 40 mM Sodium Phosphate buffer pH7.2, 20 μg/mL Zymolyase 100T (US Biological)) for 15 min at 30°C. Zymolyase was washed 3 times in spheroplasting buffer at 2,500 g and 3 times in 1X Cutsmart Buffer (NEB) at 16000 g. Cells were resuspended at a final concentration of 4.10^8^ cells/mL in 1.4 X Cutsmart buffer containing 6 pM of a long hybridization oligonucleotide (olWDH1770, **Table S2**) to restore the *Eco*RI site on the resected broken molecule, and stored at −80°C. Chromatin of 4×10^7^ cells is solubilized upon incubation at 65°C for 10 min with 0.1% SDS, and SDS is quenched by addition of 1% Triton X100. DNA is digested by 20 units of *Eco*RI-HF (NEB) at 37°C for 1 hr. Proteins are denatured by addition of 2% SDS and incubation at 55°C for 10 min. Cells are put in ice and SDS is quenched by addition of 6% Triton X100. Ligation is performed in 800 μL of a ligation mix (50 mM Tris-HCl pH 8.0, 10 mM MgCl2, 10 mM DTT, 1 mM ATP, 0.25 mg/mL BSA, 300 units of T4 DNA ligase (Bayou Biolabs)) at 16°C for 1h30. 25 μg/mL Proteinase K is added and proteins digested for 30 min at 65°C. DNA is extracted following a standard Phenol:Chloroform:Isoamyl Alcohol and isopropanol precipitation procedure. DNA pellets are re-suspended and incubated at 30°C in 50 μL 10 mM Tris HCl pH 8.0, 1 mM EDTA, 0.4 mg/mL RNAseA. The quantitative PCR was performed on a Roche LightCycler 480 machine using the SYBR Green I Master kit (Roche), according to the manufacturer instructions. After an initial denaturation phase, the cycling conditions were 95°C for 10”, 66°C for 12”, 72°C for 12”, repeated 50 times. The nature of the amplified product was confirmed by a final thermal denaturation ramp. Six reactions were performed (**Fig. S1**, primers see **Table S2**): a loading standard (*ARG4*) on which the other reactions are normalized (olWDH1760 and 1761); an intra-molecular ligation efficiency control on a 1904 bp fragment at the *DAP2* locus (olWDH1762 and 1763); a control to verify DSB formation at HOcs on Chr. V (olWDH1766 and 1767); a control of *Eco*RI restriction site digestion on the broken molecule (olWDH1768 and 1769); a control of *Eco*RI restriction digestion of a control dsDNA locus (*DAP2*; olWDH1762 and 1769); a reaction to detect the DLC chimera, *i.e.* the product of the ligation of the 5’ flanking regions of the broken molecule and the donor (olWDH1764 and 1765). DLC values were normalized on the intra-molecular ligation efficiency and in some instances corrected for the filament restriction digestion efficiency when differences exceeded ≈30%. Data were analyzed using the Light Cycler 480 Software 1.5.0.

For each mutant assayed, a wild type strain was also run in parallel to buffer for inter-experiment variations that are attributed to differences in the crosslinking efficiency. For *TOP3* and *top3-cd* overexpression experiments, an empty vector control was performed in for each experiment. In the case of the Srs2-AID degradation experiments, the control “-Auxin” treatment was run in parallel of the “+Auxin” treatment. Consequently, data are shown normalized onto parallel wild type values (for deletion mutant or point mutant), on parallel empty vector control (Top3 overexpression experiments), or on parallel DMSO-treated (Srs2-AID degradation experiments) samples.

### DLE assay

The DLE assay was performed and analyzed as described in (Piazza et al., 2018).

### Western blot

Proteins were extracted from 2.10^7^ cells by regular TCA procedure. Srs2-AID-9Myc and OsTir1-9Myc were detected with a mouse anti-c-Myc 9E11 antibody (Santa Cruz Biotechnology, sc-47694, lot F11 13) used at a 1:1000 dilution, and GAPDH was detected with a mouse anti-GAPDH GA1R from Thermo Scientific (MA5-15738, lot QG215126) at a 1:5000 dilution.

### Statistical analysis

Mutant DLC values were compared to their paired wild type or empty vector controls with a paired Student’s t-test. Other comparisons between normalized DLC values were performed using unpaired Student’s t-test. Statistical cutoff was set to α =0.05 for all tests. All statistical tests were performed under R x64 3.2.0.

### Construct sequences

The annotated sequences of the DSB-inducible and donor constructs used in this study are available as *.ape (ApE Plasmid Editor) files in **Dataset S1**.

